# A bacterial DNA repair pathway specific to a natural antibiotic

**DOI:** 10.1101/426304

**Authors:** Peter E. Burby, Lyle A. Simmons

## Abstract

All organisms possess several DNA repair pathways to maintain the integrity of their genetic material. Although there are several DNA repair pathways that are well understood, we recently identified several genes in *Bacillus subtilis* that are important for surviving treatment with drugs that damage DNA. Here, we report a drug specific DNA repair pathway in *B. subtilis*. We identified genes coding for a previously uncharacterized helicase and exonuclease, *mrfA* and *mrfB*, respectively. Deletion of *mrfA* and *mrfB* resulted in sensitivity to the DNA damaging agent mitomycin C, but not other types of DNA damage. We found that MrfAB operate independently of canonical nucleotide excision repair, forming a novel excision repair pathway in bacteria. A phylogenetic analysis demonstrates that MrfAB homologs are present in diverse bacterial phyla, and a cross-complementation assay shows that MrfAB function is conserved in closely related species. Mitomycin C is a natural antibiotic that is produced by the soil dwelling bacterium *Streptomyces lavendulae*, and *B. subtilis* is also a soil dwelling organism. The specificity of the Δ*mrfAB* phenotype suggests that MrfAB have been adapted as a countermeasure to mitomycin producing bacteria.

**Abbreviated Summary:** Bacteria possess DNA repair pathways to maintain the integrity of their genetic material. The helicase MrfA and the exonuclease MrfB are part of a mitomycin C specific DNA repair pathway in *Bacillus subtilis*. Despite being present in many bacterial species, MrfAB activity in repairing MMC damaged DNA appears to be restricted to closely related species, suggesting that these proteins have likely been adapted to the specific needs of each bacterium.

## Introduction

A defining feature of biology is the ability to reproduce, which requires replication of the genetic material. High fidelity DNA replication depends on the integrity of the template DNA which can be damaged by UV light, ionizing radiation, and numerous chemicals (Friedberg et al., 2006). Many DNA damaging agents have been used as chemotherapeutics and are also produced from natural sources such as bacteria, fungi, or plants (Demain & Vaishnav, 2011). One such naturally produced antibiotic is mitomycin C (MMC), originally isolated from *Streptomyces lavendulae* (Hata et al., 1956). MMC is produced as an inactive metabolite that must be activated by enzymatic or chemical reduction to react with DNA (Tomasz, 1995). MMC reacts specifically with guanine residues in DNA and results in three principle modifications (Bargonetti, Champeil, & Tomasz, 2010). MMC forms a mono-adduct by reacting with a single guanine, however, MMC has two reactive centers, which can result in intra-strand crosslinks on adjacent guanines on the same strand, or in inter-strand crosslinks wherein the two guanines on opposite strands of CpG sequences are covalently linked (Bizanek, McGuinness, Nakanishi, & Tomasz, 1992; Borowyborowski, Lipman, Chowdary, & Tomasz, 1990; Borowyborowski, Lipman, & Tomasz, 1990; Iyer & Szybalski, 1963; Kumar, Lipman, & Tomasz, 1992; Tomasz et al., 1986; Tomasz et al., 1987). The toxicity of these different adducts is a result of preventing DNA synthesis (Bargonetti et al., 2010).

In bacteria, MMC adducts and intra-strand crosslinks are repaired by nucleotide excision repair and inter-strand crosslinks are repaired by a combination of nucleotide excision repair and homologous recombination (Dronkert & Kanaar, 2001; Lenhart, Schroeder, Walsh, & Simmons, 2012; Noll, Mason, & Miller, 2006). Both mono-adducts and crosslinks are recognized in genomic DNA by UvrA to initiate repair (Jaciuk, Nowak, Skowronek, Tanska, & Nowotny, 2011; Kisker, Kuper, & Van Houten, 2013; Stracy et al., 2016; Weng et al., 2010). In some nucleotide excision repair models UvrB functions in complex with UvrA (Kisker et al., 2013; Truglio, Croteau, Van Houten, & Kisker, 2006; Van Houten, Croteau, DellaVecchia, Wang, & Kisker, 2005), while *in vitro* studies and a recent *in vivo* study using single molecule microscopy suggests that UvrB is recruited by UvrA (Orren & Sancar, 1989; Stracy et al., 2016). In any event, once UvrA and UvrB are present at the lesion, the subsequent step is the disassociation of UvrA and the recruitment of UvrC which incises the DNA on either side of the lesion (Orren & Sancar, 1989).

In *E. coli* there is a second UvrC like protein called Cho that can also perform the incision function (Moolenaar, van Rossum-Fikkert, van Kesteren, & Goosen, 2002; Perera, Mendenhall, Courcelle, & Courcelle, 2016). Mono-adducts and intra-strand crosslinks are removed from the DNA via UvrD helicase in *E. coli* after UvrC excision and then a DNA polymerase can resynthesize the missing nucleotides and DNA ligase can seal the gap, completing the repair process (Kisker et al., 2013; Petit & Sancar, 1999). For an inter-strand crosslink, the process requires another step because the lesion containing DNA remains covalently bonded to the opposite strand. Homologous recombination pairs the lesion containing strand with a second copy of the chromosome if present and then a subsequent round of nucleotide excision repair can remove the crosslink followed by DNA polymerase and DNA ligase to complete repair (Dronkert & Kanaar, 2001; Noll et al., 2006). Homologous recombination and UvrABC dependent nucleotide excision repair are general DNA repair pathways that participate in the repair of many different types of DNA damage.

Although the above mentioned pathways are known to function in the repair of MMC damaged DNA, it is not clear if other pathways exist in bacteria. We recently reported a forward genetic screen in *B. subtilis* where we identified two genes, *mrfA* and *mrfB* (formerly *yprA* and *yprB*, respectively) that when deleted resulted in sensitivity to MMC (Burby, Simmons, Schroeder, & Simmons, 2018). Here, we report that MrfAB are part of a MMC specific DNA repair pathway in *Bacillus subtilis*. Deletion of *mrfAB* (formerly *yprAB*) operon renders *B. subtilis* sensitive to MMC, but not to other DNA damaging agents known to be repair by canonical nucleotide excision repair. MrfAB code for a helicase and exonuclease, and we demonstrate that conserved residues required for these activities are important for their function *in vivo*. We show that MrfAB operate independent of UvrABC. We monitored DNA repair status over time using RecA-GFP as a reporter, and we show that deletion of *mrfAB* and *uvrABC* results in a synergistic decrease in RecA-GFP foci, suggesting that MrfAB are part of a novel nucleotide excision repair pathway in bacteria. We also found that MrfAB do not contribute to inter-strand crosslink repair, suggesting that MrfAB are specific to MMC mono-adducts or intra-strand crosslinks. A phylogenetic analysis shows that MrfAB homologs are present in many bacterial species and that the function of MrfAB is conserved in closely related species. Together, our study identifies a novel strategy to counteract the natural antibiotic MMC in bacteria.

## Results

### DNA damage sensitivity of Δ*mrfAB* is specific to mitomycin C

Our recent study using a forward genetic screen identified genes important for surviving exposure to DNA damage uncovering many genes that had not previously been implicated in DNA repair or the SOS-response (Burby et al., 2018). As part of this screen, we identified a gene pair, *yprAB*, in which disruption by a transposon resulted in sensitivity to MMC but not phleomycin or methyl methanesulfonate (Fig 1A) (Burby et al., 2018). Because the phenotypes are specific to MMC (see below), we rename *yprAB* to <u>m</u>itomycin <u>r</u>epair <u>f</u>actors <u>A</u> and <u>B</u> (*mrfAB*). To follow up on the phenotype of the transposon insertions we tested clean deletion strains of *mrfA* and *mrfB* and found that deletion of either gene resulted in sensitivity to MMC (Fig 1B). Further, we ectopically expressed each gene in its respective mutant background and we were capable of complementing the MMC sensitivity phenotype (Fig 1B). The absence of phenotypes with phleomycin and methyl methanesulfonate, is similar to the phenotypic profile of nucleotide excision repair (NER) mutants (Fig 1A) (Burby et al., 2018). We next asked if deletion of *mrfA* would result in sensitivity to other agents known to be repaired by NER. We tested for sensitivity to three other agents that cause DNA lesions that are repaired by NER: UV light, 4-NQO, and the DNA crosslinking agent psoralen (trioxsalen) (Petit & Sancar, 1999). Interestingly, we found that deletion of *mrfA* did not cause sensitivity to any of these agents (Fig 1C). We also tested whether the presence of *uvrAB* was masking the effect, but no additional sensitivity was observed when *mrfA* was deleted in the Δ*uvrAB* background (Fig 1C). We conclude that MrfAB are important for mitigating the toxicity of MMC-generated DNA lesions.

**Figure 1.**
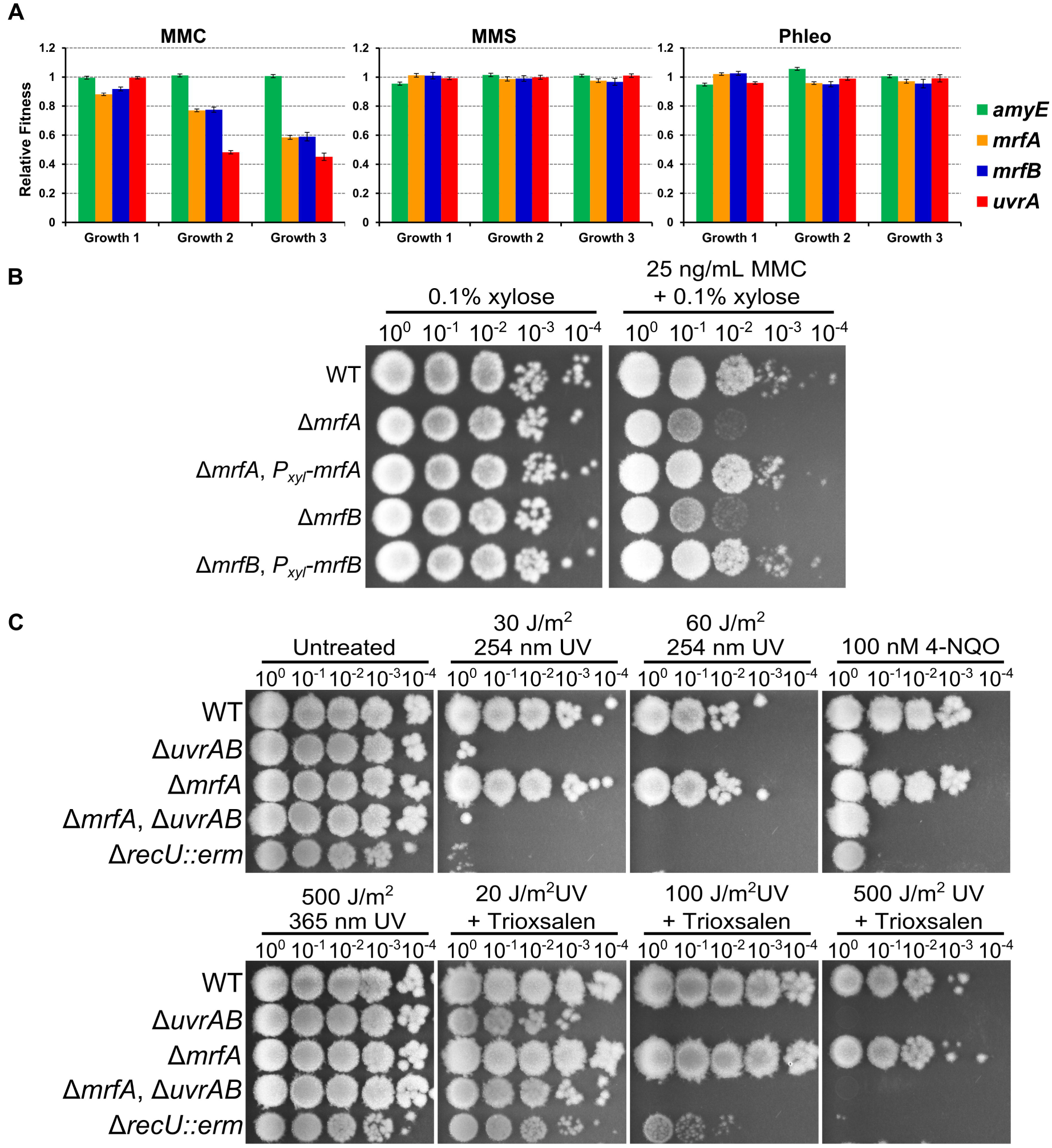
DNA damage sensitivity of Δ*mrfAB* is specific to mitomycin C. **(A)** Relative fitness plots for the indicated gene disruptions from Tn-seq experiments previously reported (Burby et al., 2018). The mean fitness is plotted as a bar graph and the error bars represent the 95% confidence interval. **(B)** Spot titer assay using strains with the indicated genotypes grown on the indicated media. **(C)** Spot titer assay using strains with the indicated genotypes grown on the indicated media. For UV treatments, cells were exposed to the indicated dose after serial dilutions were spotted on the media. For trioxsalen plates 1 μg/mL was used and the UV wavelength for irradiation was 365 nm.

### MrfA and MrfB function in the same pathway

The phenotypes of *mrfA* and *mrfB* mutants were identical (Fig 1A&B), and the two genes are predicted to be an operon. We hypothesized that MrfA and MrfB function together. We tested this hypothesis using an epistasis analysis. We found that deletion of both genes gave the same sensitivity to MMC as each single mutant (Fig 2A), indicating that they function in the same pathway. If MrfAB function together, we wondered whether these proteins interact. We employed a bacterial two-hybrid assay, and found that MrfA and MrfB formed a robust interaction (Fig 2B). Next, we asked if we could localize the interaction to a particular domain.

**Figure 2.**
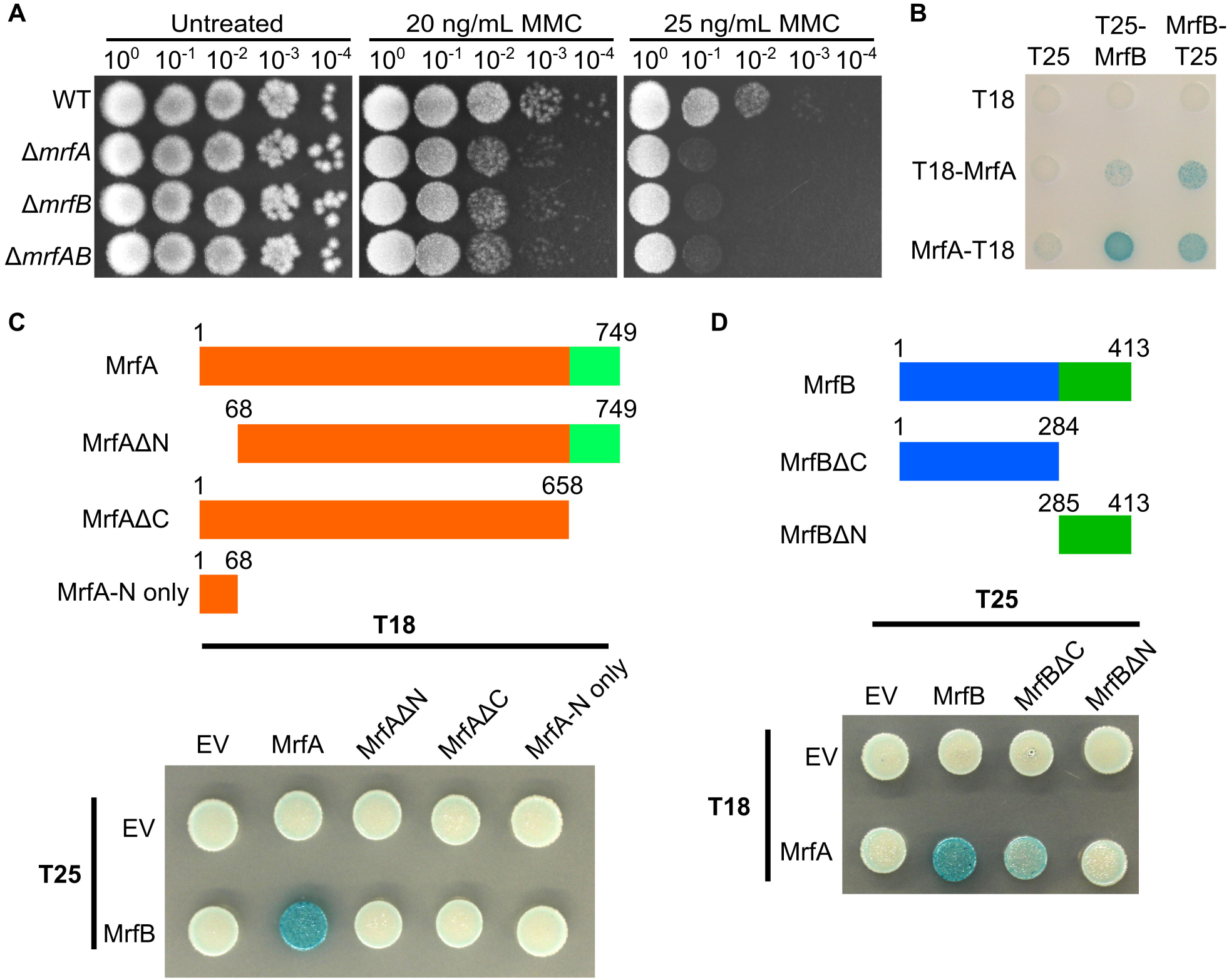
MrfA and MrfB function in the same pathway. **(A)** Spot titer assay using strains with the indicated genotypes grown on the indicated media. **(B)** Bacterial two-hybrid assay using the indicated T18 and T25 fusions. **(C)** MrfA constructs used in deletion analysis of MrfA-MrfB interaction (upper) and a bacterial two-hybrid assay using T25-MrfB and the indicated MrfA-T18 fusions (lower). **(D)** MrfB constructs used in deletion analysis of MrfA-MrfB interaction (upper) and a bacterial two-hybrid assay using MrfA-T18 and the indicated T25-MrfB fusions (lower).

We performed a deletion analysis with MrfA and found that deletion of either the N-terminus or the C-terminus was sufficient to abolish the interaction with MrfB (Fig 2C), and the N-terminus of MrfA was not sufficient for MrfB interaction (Fig 2C). We tested whether the N-terminus or C-terminus of MrfB was required for MrfA interaction. We found that the C-terminus of MrfB was not required, though the signal was reduced, whereas deletion of the N-terminus of MrfB abolished the interaction with MrfA (Fig 2D). We conclude that MrfAB interaction is specific and that these proteins function together in mitigating MMC toxicity.

### MrfA helicase motifs and C-terminus is required for function *in vivo*

MrfA is a predicted DEXH box helicase containing a C-terminal domain of unknown function (Fig S1 and S2A). The C-terminal domain of unknown function contains four conserved cysteines that are thought to function in coordinating a metal ion (Shi et al., 2011; Yakovleva & Shuman, 2012). We initially searched for a similar helicase in other well studied organisms. We were unable to identify a clear homolog of MrfA in *E. coli*, however, Hrq1 from *Saccharomyces cerevisiae* shares the same domain structure with 32% identity and 55% positives. Hrq1 has been shown to be a RecQ family helicase with 3□ → 5□ helicase activity and has been observed to exist as a heptamer (Bochman, Paeschke, Chan, & Zakian, 2014; Rogers et al., 2017). We performed an alignment with Hrq1 and identified helicase motifs typical of super family 2 helicases (Fig S1). A homolog of MrfA from *Mycobacterium smegmatis* has also been shown to be a 3□ → 5□ helicase, however, unlike Hrq1 SftH exists as a monomer in solution (Yakovleva & Shuman, 2012). To address whether MrfA helicase activity was important for function we used a complementation assay using variants containing alanine substitutions in several conserved helicase motifs. Mutations in helicase motif I (K82A), motif II (DE185-186AA), and motif III (S222A) all failed to complement a *mrfA* deficiency (Fig S2B). Intriguingly, when motif Ib (T134V) was mutated *mrfA* MMC sensitivity could still be complemented (Fig S2B). We asked whether the C-terminal domain of unknown function and the conserved cysteines were required for function. Deletion of the entire C-terminal domain, mutation of the first two cysteines, or mutation of all four cysteines all resulted in a failure to complement MMC sensitivity in a Δ*mrfA* strain (Fig S2B). Together our data suggest that both helicase activity and the C-terminal domain are required for MrfA function *in vivo*.

### MrfB is a metal dependent exonuclease

MrfB is predicted to be a DnaQ-like exonuclease and to have a three tetratrichopeptide repeats at its C-terminus (Fig 3A). To search for putative catalytic residues in MrfB, we aligned MrfB to ExoI, ExoX, and DnaQ from *E. coli* (Fig S3A). MrfB has the four acidic residues typical of DnaQ like exonucleases (Fig S3A). This type of nuclease also has a histidine located proximal to the last aspartate (Yang, 2011), and we identified two histidines, one of which was conserved (Fig S3A, conserved histidine highlighted in red and the other in green). DnaQ exonucleases coordinate a metal ion that is used in catalysis (Yang, 2011). We hypothesized that MrfB catalytic residues would cluster together in the tertiary structure. We modelled MrfB using Phyre2.0 (Kelley, Mezulis, Yates, Wass, & Sternberg, 2015), which used DNA polymerase epsilon catalytic subunit A (pdb structure c5okiA (Grabarczyk, Silkenat, & Kisker, 2018)), and found that the conserved aspartates and glutamate indeed clustered together in the model (Fig S3B).

**Figure 3.**
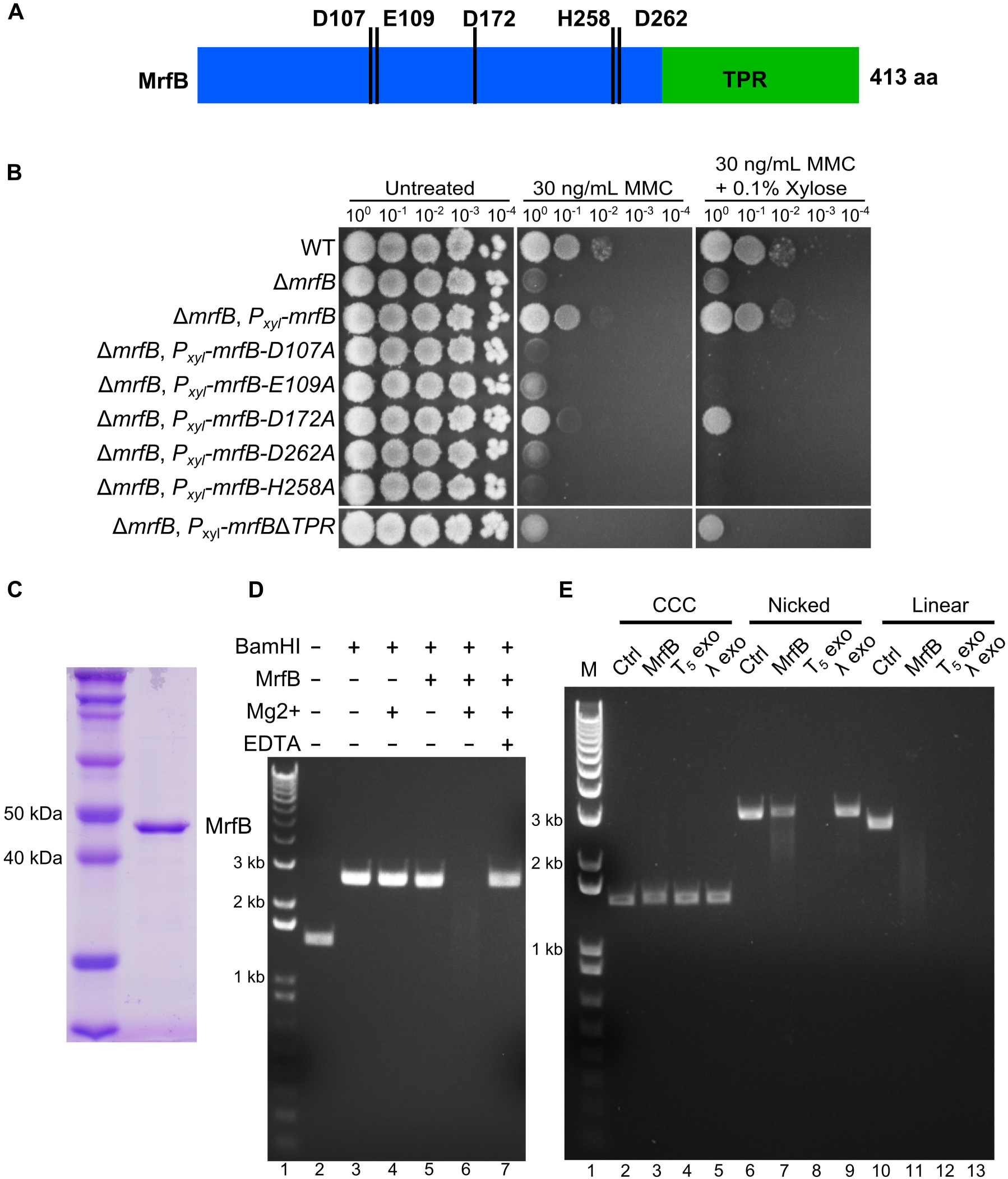
MrfB is a metal-dependent exonuclease. **(A)** A schematic of MrfB depicting putative catalytic residues and C-terminal tetra-trichopeptide repeat (TPR) domain. **(B)** Spot titer assay using strains with the indicated genotypes spotted on the indicated media. **(C)** Purified MrfB stained with Coomassie blue. **(D)** Exonuclease assay using pUC19 linearized with BamHI (lanes 3-7). Reactions were incubated at 37°C for 15 minutes with or without MrfB, MgCl_2_, or EDTA as indicated, and separated on an agarose gel stained with ethidium bromide. Lane 1 is a 1 kb plus molecular weight marker (M) and lane 2 is undigested pUC19 plasmid. **(E)** Exonuclease assay testing substrate preference. The indicated exonucleases were incubated with a closed covalent circular plasmid (CCC), a nicked plasmid (Nicked) or a linear plasmid (Linear) in the presence of Mg^2+^ at 37°C for 10 minutes. Reaction products were separated on an agarose gel stained with ethidium bromide. Lane 1 is a 1 kb plus molecular weight marker (M).

Interestingly, we found that the histidine conserved in the *E. coli* exonucleases was facing the opposite direction, whereas the non-conserved histidine was facing the putative catalytic residues in the MrfB model (Fig S3C). An alignment of MrfB with homologs demonstrates that the histidine (labeled in green) facing the other putative catalytic residues is conserved in putative MrfB homologs, whereas the other is not (see supplemental text). To test whether these residues were important for function, we used variants with alanine substitutions at each putative catalytic residue in a complementation assay. We found that all five mutants could not complement the Δ*mrfB* mutant (Fig 3B).

With these results we wanted to test whether MrfB had exonuclease activity *in vitro*. We expressed and purified MrfB to homogeneity as determined by SDS-PAGE (Fig 3C). We tested for exonuclease activity using a plasmid linearized by restriction digest. We found that MrfB could degrade linear dsDNA and that Mg^2+^ was required, demonstrating that MrfB is a metal-dependent exonuclease (Fig 3D). With exonuclease activity established we tested the substrate preference of MrfB using a closed circular covalent plasmid (CCC), a nicked plasmid or a linear plasmid using T_5_ and *λ* exonucleases as controls. T_5_ exonuclease is able to degrade both nicked and linear substrates but cannot degrade a super-coiled plasmid (Sayers & Eckstein, 1990, 1991). In contrast, λ exonuclease can only degrade a linear substrate (Little, 1981). The T_5_ and lambda exonuclease controls performed as predicted, and MrfB demonstrated robust activity using a linear substrate and very slight activity using a nicked substrate (Fig 3E). We conclude that MrfB is a metal-dependent exonuclease with a preference for linear DNA.

### MrfAB function independent of UvrABC dependent nucleotide excision repair

Given that DNA damage sensitivity in *mrfAB* mutants was restricted to MMC and that both proteins have nucleic acid processing activities, we hypothesized that MrfAB were part of a nucleotide excision repair pathway. We tested whether MrfAB were within the canonical, UvrABC dependent nucleotide excision repair pathway using an epistasis analysis. We found that deletion of *mrfA* or *mrfB* rendered *B. subtilis* hypersensitive to MMC in the absence of *uvrAB* (Fig 4A), *uvrC*, or *uvrABC* (Fig 4B). We also verified that *uvrABC* indeed function as a single pathway using an epistasis analysis (Fig S4), which differs from *E. coli* (Lage, Goncalves, Souza, de Padula, & Leitao, 2010; Perera et al., 2016). To test whether deletion of *mrfAB* have an effect on acute treatment with MMC, we performed an epistasis analysis using a MMC survival assay. We tested mutants in *mrfAB*, *uvrABC*, and the double pathway mutant. We found that deletion of *mrfAB* had a limited effect on acute sensitivity to MMC whereas deletion of *uvrABC* had a significant decrease in survival following MMC treatment (Fig 4C). Deletion of both pathways resulted in hypersensitivity to acute MMC exposure, suggesting that MrfAB are part of a second nucleotide excision repair pathway. We conclude that MrfAB and UvrABC are part of two distinct pathways for MMC repair.

**Figure 4.**
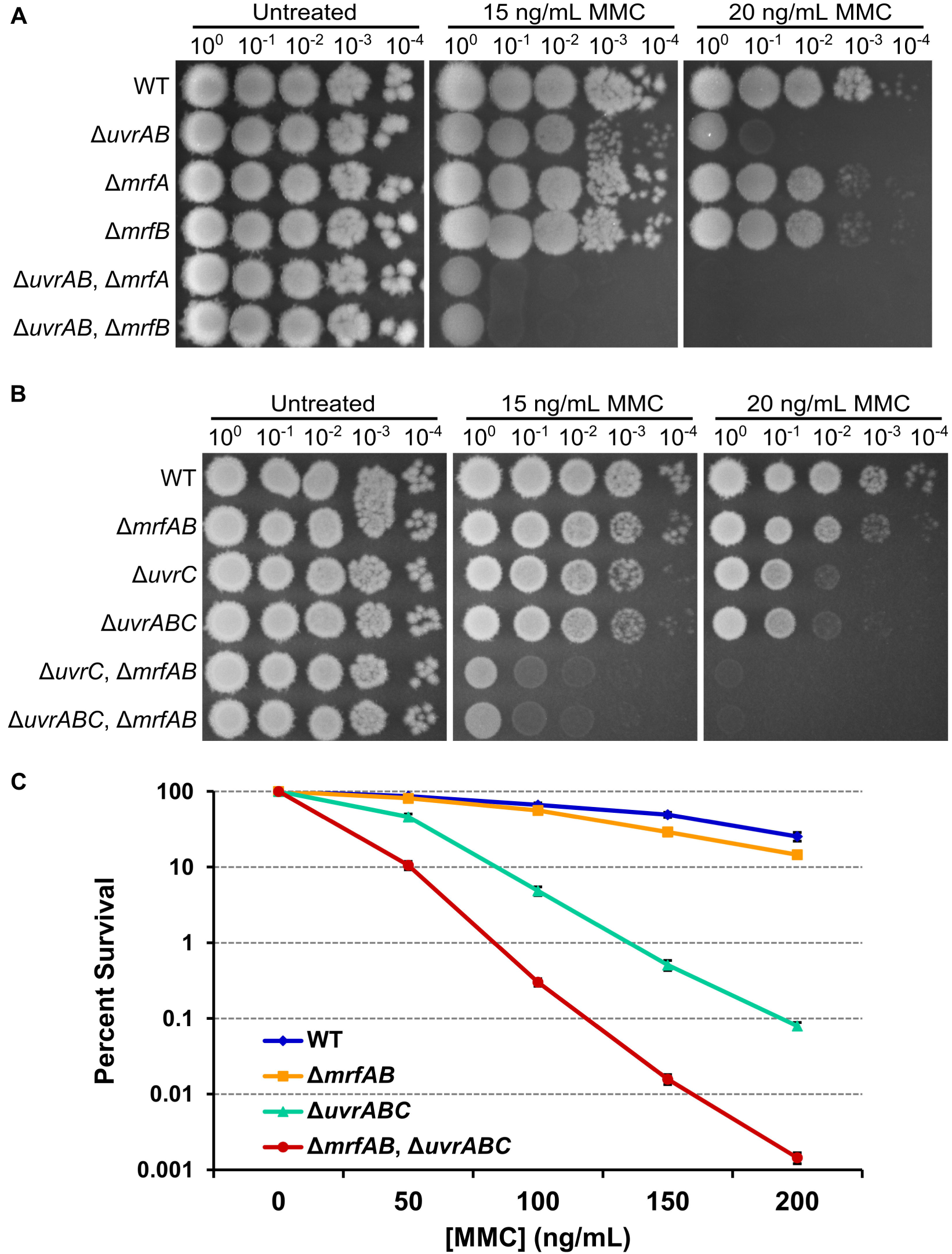
MrfAB function independent of UvrABC dependent nucleotide excision repair. **(A & B)** Spot titer assays using strains with the indicated genotypes grown on the indicated media. **(C)** Survival assay using strains with the indicated genotypes. The y-axis is the percent survival relative to the untreated (0 ng/mL) condition. The x-axis indicates the concentration of MMC used for a 30 minute acute exposure. The data points represent the mean of three independent experiments performed in triplicate (n=9) ± SEM.

### MrfAB are not required for unhooking inter-strand DNA crosslinks

The synergistic sensitivity to MMC observed in the double pathway mutant suggests that MrfAB are part of a novel nucleotide excision repair pathway. Previous studies have demonstrated that RecA-GFP forms foci in response to DNA damage such as treatment with MMC (Kidane & Graumann, 2005; Simmons et al., 2009; Simmons, Grossman, & Walker, 2007). Therefore, to test whether DNA repair was affected by the absence of *mrfAB*, *uvrABC*, or both pathways, we used a RecA-GFP fusion to monitor DNA repair status over time following treatment with MMC (Fig 5A & S5). We quantified the percentage of cells containing a focus or foci of RecA-GFP, and found an increase in RecA-GFP focus formation over time (Fig 5B). In all three mutant strains there was a significant increase in RecA-GFP foci prior to MMC addition (Fig 5B). We found that deletion of *mrfAB* did not have a significant impact on RecA-GFP focus formation (Fig 5B). Deletion of *uvrABC* led to a slight decrease in RecA-GFP focus formation (Fig S5).

**Figure 5.**
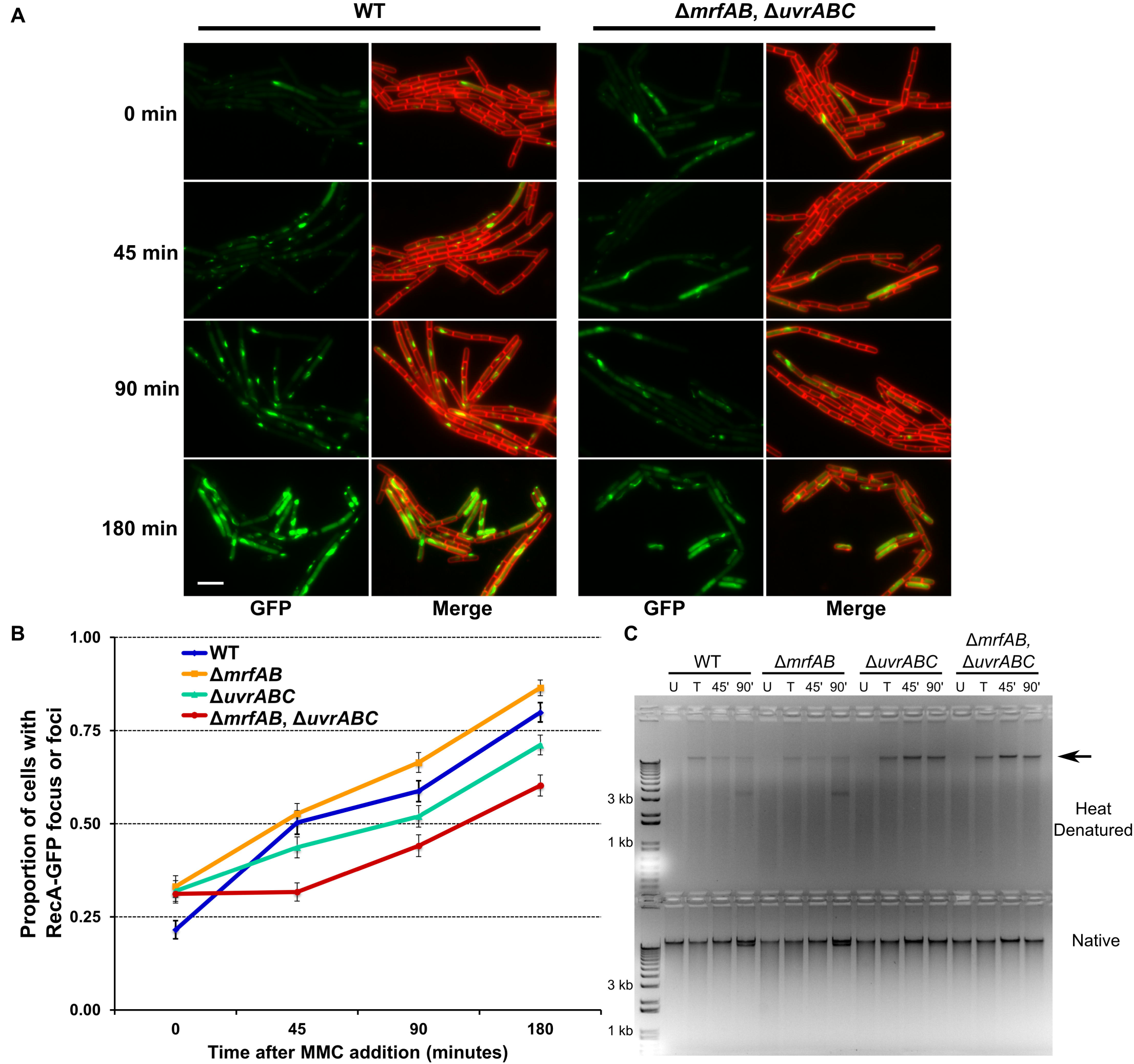
MrfAB are not required for unhooking inter-strand DNA crosslinks. **(A)** Representative micrographs of strains containing RecA-GFP expressed from the native locus in addition to the indicated genotypes. Images were captured at the indicated times following MMC addition (5 ng/mL). RecA-GFP is shown in green and the merged images show RecA-GFP (green) and membranes stained with FM4-64 (red). The white bar indicates 5 µm **(B)** Percentage of cells with a RecA-GFP focus or foci over the indicated time course of MMC treatment (5 ng/mL). The error bars represent the 95% confidence interval **(C)** DNA crosslinking and repair assay. Chromosomal DNA from untreated samples (U), 1 μg/mL MMC treated samples (T), and recovery samples (45’ and 90’) were heat denatured and snap cooled (upper) or native chromosomal DNA (lower) was separated on an agarose gel stained with ethidium bromide. A 1 kb plus molecular weight marker is shown in the first lane.

The double pathway mutant had a significant decrease in RecA-GFP foci (Fig 5B). With these results we suggest that RecA is responding to excision repair gaps that occur after removal of the MMC adduct. These results further support the conclusion that MrfAB participate in the repair of MMC damaged DNA.

As stated previously, MMC results in several DNA lesions, one of which is the inter-strand crosslink. We asked whether one or both pathways contribute to unhooking DNA crosslinks *in vivo*. To address this question we treated *B. subtilis* strains with MMC and then allowed them to recover for 45 or 90 minutes and monitored DNA crosslinks by denaturing and snap cooling the DNA, which will allow for renaturing of crosslinked DNA but not non-crosslinked DNA (Iyer & Szybalski, 1963). We found that in WT and Δ*mrfAB* cells we could detect some crosslinked DNA that decreased slightly over time (Fig 5C). In the absence of *uvrABC* there was a significant stabilization of crosslinked DNA that did not decrease over time and deleting *mrfAB* had no effect in the *uvrABC* mutant on crosslink stabilization (Fig 5C). We conclude that UvrABC are the primary proteins responsible for repair of inter-strand crosslinks and MrfAB likely repair the more abundant mono-adducts (Warren, Maccubbin, & Hamilton, 1998) and potentially intra-strand crosslinks that form.

### MrfAB are conserved in diverse bacterial phyla

Given the specificity of MrfAB for MMC, we wondered how conserved *mrfA* and *mrfB* are in bacteria. We performed a PSI-BLAST search using MrfA or MrfB against the proteomes of bacterial organisms from several phyla (Fig 6A; Table S4). We found that MrfA and MrfB are both present in organisms from 5 different phyla, though MrfA is more broadly conserved in bacteria (Fig 6A). To test if MrfA and MrfB function is conserved, we attempted to complement MMC sensitivity using codon-optimized versions of the homologs from three organisms, *Bacillus cereus*, *Streptococcus pneumoniae*, and *Pseudomonas aeruginosa*. We found that expression of *Bc*-*mrfA* and *Bc*-*mrfB* were capable of complementing the respective deletion (Fig 6B). Interestingly, *Sp*-*mrfB* complemented, but *Sp*-*mrfA* did not (Fig 6B). The more distantly related homologs from *P. aeruginosa* were not able to complement the corresponding deletion alleles (Fig 6B). We conclude that MrfA and MrfB function is conserved in closely related species, and that they likely have been adapted to other uses in more distantly related bacteria.

**Figure 6.**
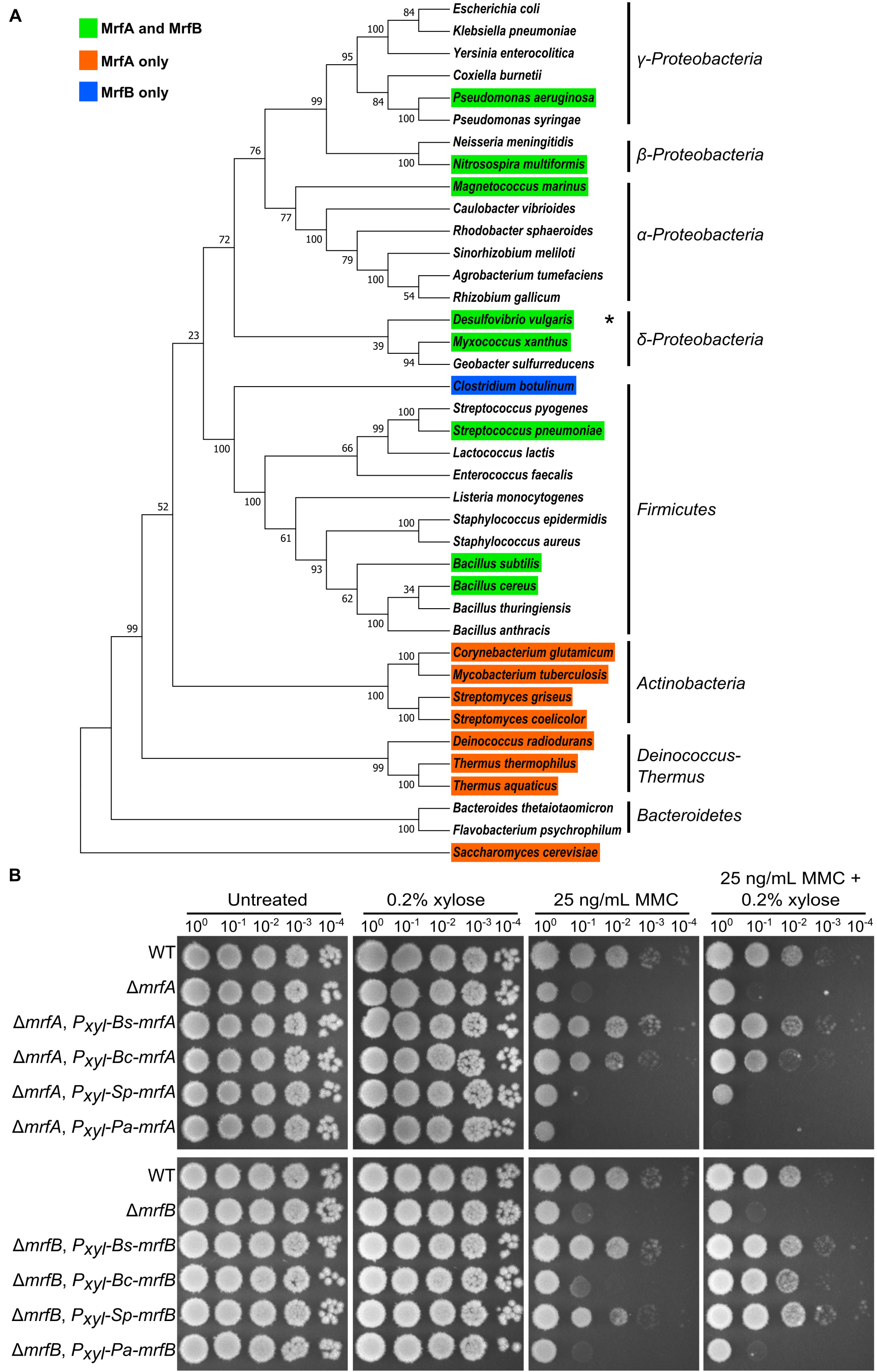
MrfAB are conserved in diverse bacterial *phyla*. **(A)** A rooted phylogenetic tree constructed using 16s rRNA sequences (18s rRNA for *S. cerevisiae*), aligned with muscle (Edgar, 2004), using the neighbor joining method (Saitou & Nei, 1987), and the evolutionary distances were calculated using the p-distance method (Nei & Kumar, 2000). The percentage of replicate trees that resulted in the associated species clustering together in a bootstrap test (500 replicates) is indicated next to the branches (Felsenstein, 1985). Evolutionary analysis was performed in MEGA (Kumar, Stecher, & Tamura, 2016). ^∗^In this organism MrfA and MrfB homologs are fused into a single protein. **(B)** Spot titer assay using codon optimized versions of MrfA and MrfB from the indicated species to complement Δ *mrfA* or Δ *mrfB* (lower).

## Discussion

MrfAB are founding members of a novel bacterial nucleotide excision repair pathway. From our results, the current model is that MrfA recognizes a MMC mono-adduct or intra-strand crosslink and uses its helicase activity to separate the MMC containing strand, while MrfB degrades the displaced DNA strand ultimately removing the lesion (Fig 7). The initial finding that sensitivity to DNA damage in *mrfAB* mutants is specific to MMC suggested a drug specific repair pathway. The observation that RecA-GFP foci changes in a synergistic manner with deletion of both *uvrABC* and *mrfAB* suggests that MrfAB are acting as a second excision repair pathway. The major source of toxicity from MMC has long been thought to be the inter-strand crosslink (Bargonetti et al., 2010). We found that MrfAB do not contribute to inter-strand crosslink repair, and yet deletion of *mrfAB* in the *uvrABC* mutant resulted in a significant decrease in survival following MMC treatment. These observations strongly suggest that the mono-adducts and/or the intra-strand crosslink make a significant contribution to the overall toxicity of MMC. Therefore, through identifying a new repair pathway in bacteria, we are able to shed new light on the toxicity profile of a well-studied, natural antibiotic.

**Figure 7.**
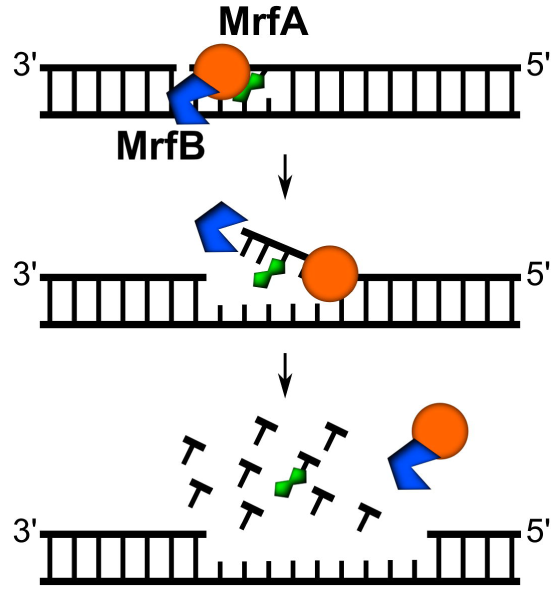
A model for MrfAB mediated nucleotide excision repair. MrfA recognizes an MMC adduct and recruits MrfB. MrfA uses its helicase activity to separate the strand containing the MMC adduct, facilitating MrfB-dependent degradation of the adduct containing DNA.

The specificity of the Δ*mrfAB* phenotype suggests that lesion recognition depends on MMC adduct structure. Our previously reported forward genetic screen did not identify other candidates for this pathway (Burby et al., 2018). Thus, we hypothesize that lesion recognition is a function carried out by either MrfA, MrfB, or by both proteins in complex. MrfA is a helicase with a C-terminal domain of unknown function containing four well conserved cysteines. A high throughput X-ray absorption spectroscopy study of over 3000 proteins including MrfA reported finding that MrfA binds to zinc (Shi et al., 2011). Intriguingly, UvrA, the recognition factor of canonical nucleotide excision repair, also contains a zinc finger which is required for regulating recognition of damaged DNA (Croteau et al., 2006). Indeed, three of the four recognition factors in eukaryotic nucleotide excision repair, XPA, RPA, and TFIIH also each contain a zinc finger component (Petit & Sancar, 1999). Therefore, it is tempting to speculate that MrfA functions as the lesion recognition factor through its putative C-terminal zinc finger domain.

MrfAB have likely been adapted to specific needs in different bacteria. We speculate that MrfAB specificity for MMC is a reflection of habitat overlap between *B. subtilis* and mitomycin producing bacteria such as *S. lavendulae*. Thus, MrfAB were adapted to effectively help *B. subtilis* compete in habitats where MMC is produced. Given that only closely related species could substitute for MrfA and MrfB function in *B. subtilis*, we hypothesize that the MMC specific repair activity is restricted to those species. In fact, the homologs present in *P. aeruginosa* have diverged significantly (Table S4). The N-terminus of *Pa*-MrfA is quite different from that of *Bs*-MrfA, and the C-terminal TPR domain of MrfB is completely absent in *Pa*-MrfB (see supplemental alignments), consistent with the notion that MrfAB function has diverged in more distantly related bacteria. We recently investigated the mismatch repair homolog MutS2 and arrived at a similar conclusion—MutS2 has been adapted to the specific DNA repair needs of different organisms. MutS2 in *B. subtilis* promotes homologous recombination (Burby & Simmons, 2017), whereas MutS2 in several other organisms inhibits homologous recombination (Damke, Dhanaraju, Marsin, Radicella, & Rao, 2015; Fukui et al., 2008; Pinto et al., 2005; Wang & Maier, 2017). The reality that distantly related organisms have adapted their genetic repertoire inherited from the most recent common ancestor would seem obvious. Still, a major thrust of biological exploration is often to examine processes that are highly conserved. While well conserved processes are often critical for more organisms, it is the divergent functions that make each organism unique, which is a property of inherent value found throughout nature.

## Materials and Methods

### Bacteriological methods

All *B. subtilis* strains used in this study are isogenic derivatives of PY79 (Youngman, Perkins, & Losick, 1984), and listed in Table S1. Detailed construction of strains and plasmids and oligonucleotides used in this study are described in the supplemental text. Plasmids and oligonucleotides are listed in Supplemental Tables S2 and S3, respectively. Media used to culture *B. subtilis* include LB (10 g/L NaCl, 10 g/L tryptone, and 5 g/L yeast extract) and S7_50_ minimal media with 2% glucose (1× S7_50_ salts (diluted from 10× S7_50_ salts: 104.7g/L MOPS, 13.2 g/L, ammonium sulfate, 6.8 g/L monobasic potassium phosphate, pH 7.0 adjusted with potassium hydroxide), 1× metals (diluted from 100× metals: 0.2 M MgCl_2_, 70 mM CaCl_2_, 5 mM MnCl_2_, 0.1 mM ZnCl_2_, 100 μg/mL thiamine-HCl, 2 mM HCl, 0.5 mM FeCl_3_), 0.1% potassium glutamate, 2% glucose, 40 μg/mL phenylalanine, 40 μg/mL tryptophan). Selection of *B. subtilis* strains was done using spectinomycin (100 μg/mL) or chloramphenicol (5μg/mL).

### Spot titer and survival assays

Spot titer assays were performed as described previously (Burby et al., 2018). Survival assays were performed as previously described (Burby et al., 2018), except cells were treated at a density of OD_600_ = 1 instead of 0.5.

### Microscopy

Strains containing RecA-GFP were grown on LB agar + 100 μg/mL spectinomycin at 30°C overnight. Plates were washed with S7_50_ minimal media with 2% glucose. Cultures of S7_50_ minimal media with 2% glucose and 100 μg/mL spectinomycin were inoculated at an OD_600_ = 0.1 and incubated at 30°C protected from light until an OD_600_ of about 0.3 (about 3.5 hours). Cultures were treated with 5 ng/mL MMC and samples were taken for imaging prior to MMC addition, 45 minutes, 90 minutes, and 180 minutes after MMC addition. The vital membrane stain FM4-64 was added to 2 μg/mL and left at room temperature for five minutes. Samples were transferred to 1% agarose pads containing 1× Spizizen salts as previously described (Burby et al., 2018). Images were captured using an Olympus BX61 microscope using 250 ms exposure times for both FM4-64 (membranes) and GFP. RecA-GFP foci were determined by using the find maxima function in ImageJ with the threshold set to the background of the image by looking at a line trace of an area without cells. The number of cells with foci was determined by taking the total number of foci and subtracting the foci greater than one in cells having multiple foci (*i.e.*, if a cell had two foci, one would be subtracted and if a cell had 3 foci two would be subtracted and so on). The percentage was determined by dividing the number of cells with a focus or foci by the total number of cells observed.

### DNA crosslinking assay

Strains of *B. subtilis* were struck out on LB agar and incubated at 30°C overnight. Plates were washed with LB and samples of 0.5 mL OD_600_ = 3 were aliquoted. One sample was untreated and three samples were treated with 1 μg/mL MMC. Samples were incubated at 37°C for 1 hour. For the untreated and MMC treatment samples, one volume (0.5 mL) of methanol was added and samples were mixed by inversion. Samples were harvested via centrifugation (12,000 *g* for 5 minutes, washed twice with 0.5 mL 1× PBS pH 7.4 and stored at −20°C overnight). For recovery samples, cells from the remaining two treated samples were pelleted via centrifugation (10,000 *g* for 5 minutes) washed twice with 1 mL LB media and then re-suspended in 0.6 mL LB media. Samples were then transferred to 14 mL round bottom culture tubes and incubated at 37°C on a rolling rack for 45 or 90 minutes. An equal volume (0.6 mL) of methanol was added and samples were mixed by inversion. Samples were harvested as stated above and stored at −20°C overnight. Chromosomal DNA was extracted using a silica spin-column as previously described (Burby et al., 2018). Samples were normalized by A_260_ to 15 ng/μL. Samples were heat denatured by incubating at 100°C for 6 minutes followed by placing directly into an ice-water bath for 5 minutes. For native samples and heat denatured samples, 300 ng and 600 ng, respectively, were loaded onto a 0.8% agarose gel with ethidium bromide and electrophoresed at 90 volts for approximately one hour.

### Bacterial two-hybrid assays

Bacterial two-hybrid assays were performed as described (Burby et al., 2018; Karimova, Gauliard, Davi, Ouellette, & Ladant, 2017)

### MrfB protein purification

MrfB was purified from *E. coli* cells as follows. 10×His-Smt3-MrfB was expressed from plasmid pPB97 in *E. coli* NiCo21 cells (NEB) at 37°C. Cells were pelleted and resuspended in lysis buffer (50 mM Tris pH7.5, 300 mM NaCl, 5% sucrose, 25 mM imidazole, 1× Roche protease inhibitor cocktail). Cells were lysed via sonication and lysates were clarified via centrifugation: 18,000 rpm (Sorvall SS-34 rotor) for 45 minutes at 4°C. Clarified lysates were loaded onto Ni^2+^-NTA-agarose pre-equilibrated in lysis buffer in a gravity column. The column was washed with 25 column volumes wash buffer (50 mM Tris pH 7.5, 500 mM NaCl, 10% (v/v) glycerol, 40 mM imidazole). MrfB was eluted from the column by cleavage of the 10×His-Smt3 tag using 6×His-Ulp1 in 10 column volumes of digestion buffer (50 mM Tris pH 7.5, 150 mM NaCl, 10% glycerol, 10 mM imidazole, 1 mM DTT, and 20 μg/mL 6×His-Ulp1) at room temperature for 150 minutes. The eluate containing untagged MrfB was collected as the flow through. MrfB was concentrated using a 10 kDa Amicon centrifugal filter. MrfB was loaded onto a HiLoad superdex 200-PG 16/60 column pre-equilibrated with gel filtration buffer (50 mM Tris pH 7.5, 250 mM NaCl, and 5% (v/v) glycerol). The column was eluted with gel filtration buffer at a flow rate of 1 mL/min. Peak fractions were pooled, glycerol was added to a final concentration of 20%, and concentrated using a 10 kDa Amicon centrifugal filter. MrfB aliquots were frozen at a final concentration of 2.6 μM in liquid nitrogen, and stored at 80°C.

### Exonuclease assays

Exonuclease reactions (20 μL) were performed in 25 mM Tris pH 7.5, 20 mM KCl, and 5 mM MgCl_2_ as indicated in the figure legends. The plasmid pUC19 was used as a substrate at a concentration of 13.5 ng/μL. To generate linear or nicked substrate, pUC19 was first incubated with BamHI-HF (NEB) or Nt.BSPQ1 (NEB), respectively, for 30 minutes at 37°C. To test metal dependency of MrfB, the linearized pUC19 was purified using a silica spin-column. Reactions were initiated by adding MrfB to 130 nM, 10 units of T5 exonuclease (NEB), or 5 units of *λ* exonuclease (NEB) and incubating at 37°C as indicated in the figure legends. Reactions were terminated by the addition of 8 μL of nuclease stop buffer (50% glycerol and 100 mM EDTA) and reaction products were separated by agarose gel electrophoresis.

### Phylogenetic analysis

The protein sequences of MrfA (AHA78094.1) and MrfB (AHA78093.1) were used in a PSI-BLAST search in the organisms listed in Table S4. If a putative homolog was detected, the coverage and percent identity were both recorded (Table S4). For MrfA, the protein was called a homolog if the DEXH helicase domain, the C-terminal domain, and the four conserved cysteines were all present. For MrfB, the protein was called a homolog if the putative catalytic residues were conserved.

## Acknowledgements

PEB was supported by a predoctoral fellowship #DGE1256260 from the National Science Foundation. National Institutes of Health grant R01 GM107312 to LAS supported this work.

## Author contributions

This study was conceived and designed by P.E.B. and L.A.S. Experiments were performed by P.E.B. Data analysis was performed by P.E.B and L.A.S. The manuscript was written and revised by P.E.B. and L.A.S.

